# Nanomolar range of FAM237B can activate receptor GPR83

**DOI:** 10.1101/2023.05.04.539513

**Authors:** Hao-Zheng Li, Ya-Fen Wang, Wen-Feng Hu, Ya-Li Liu, Zeng-Guang Xu, Zhan-Yun Guo

**Affiliations:** Research Center for Translational Medicine at East Hospital, School of Life Sciences and Technology, Tongji University, Shanghai, China

**Keywords:** FAM237A, FAM237B, GPR83, Agonist, Binding

## Abstract

The orphan G protein-coupled receptor 83 (GPR83) is implicated in the regulation of energy metabolism and certain anxiety-related behaviors. Our recent study confirmed that family with sequence similarity 237 member A (FAM237A), also known as neurosecretory protein GL (NPGL), is an efficient agonist for GPR83, but did not support the proprotein convertase subtilisin/kexin type 1 inhibitor (PCSK1N, also known as proSAAS)-derived peptide PEN and the procholecystokinin-derived peptide proCCK56-63 as ligands of this receptor. FAM237B (also known as NPGM) is a paralog of FAM237A that was previously reported as a weak agonist for GPR83 with approximately 100-fold lower activity in an inositol 1-phosphate accumulation assay. In the present study, we prepared mature human FAM237B via an intein-fusion approach and measured its activity towards human GPR83 via a NanoLuc Binary Technology (NanoBiT)-based ligand□receptor binding assay and a NanoBiT-based β-arrestin recruitment assay. Mature FAM237B displayed moderately lower activity than its paralog FAM237A in these binding and activation assays, but could cause a significant activation effect at the nanomolar range (1□10 nM). Thus, FAM237B appears to be another endogenous agonist for receptor GPR83.

## Introduction

Currently, the A-class G protein-coupled receptor 83 (GPR83) is officially classified as an orphan receptor. It is widely present in vertebrates and is highly conserved from fishes to mammals. GPR83 is primarily expressed in the brain, and is involved in the regulation of energy metabolism and certain anxiety-related behaviors (Dubins et al. 2012; Fakira et al. 2019, 2021; Müller et al. 2013). The human *GPR83* gene can produce two transcripts (NM_016540 and NM_001330345) that either encode longer isoform 1 with seven transmembrane domains (TMDs) or shorter isoform 2 lacking TMD3. In previous publications and in the present study, GPR83 refers to the longer isoform 1 unless otherwise stated.

In recent years, the proprotein convertase subtilisin/kexin type 1 inhibitor (PCSK1N, also known as proSAAS)-derived peptide PEN (Gomes et al. 2016), the procholecystokinin (proCCK)-derived peptide proCCK56-63 (Mack et al. 2022), and family with sequence similarity 237 member A (FAM237A) (Sallee et al. 2020) were all reported as efficient agonists for GPR83. In a recent study (Li et al. 2023), we confirmed that human FAM237A binds to human GPR83 with nanomolar range affinity and activates it with nanomolar range efficiency using the NanoLuc Binary Technology (NanoBiT)-based ligand−receptor binding assay, fluorescent ligand-based visualization, and the NanoBiT-based β-arrestin recruitment assay; however, we did not detect any interaction between peptide PEN and proCCK56-63 with this receptor using these assays. Thus, it seemed that FAM237A, rather than peptide PEN and proCCK56-63, is the ligand for GPR83.

FAM237A was also named neurosecretory protein GL (NPGL) by Ukena’s group (Ukena et al. 2014), and its biological function has been studied by this group in recent years (Fukumura et al. 2021a, 2021b, 2021c; Iwakoshi-Ukena et al. 2017; Matsuura et al. 2017; Narimatsu et al. 2021, 2022a, 2022b, 2022c; Shikano et al. 2018a, 2019, 2020). Administration or overexpression of FAM237A in murine or avian models typically increased food intake and fat accumulation, suggesting this neuropeptide is implicated in the regulation of energy metabolism. Consistently, knockout of *Gpr83* in a mouse model led to a leaner phenotype with less fat accumulation (Müller et al. 2013). Thus, it seemed that FAM237A is an endogenous agonist for receptor GPR83.

FAM237B (named NPGM by Ukena’s group) is a paralog of FAM237A. FAM237B is also widely present from fish to mammals and is conserved in evolution (Fig. S1). Both FAM237B and FAM237A are synthesized *in vivo* as precursors, including an N-terminal signal peptide, a mature peptide, and a C-terminal propeptide. At the conjunction of their predicted mature peptide and propeptide, a Gly residue and a dibasic cleavage site are absolutely conserved (Fig. S1), implying C-terminal α-amidation of their mature peptide. Moreover, both proteins contain two absolutely conserved Cys residues in their predicted mature peptide (Fig. S1), implying the formation of an intramolecular disulfide bond. Preliminary studies suggested that the biological function of FAM237B is similar to that of FAM237A (Kato et al. 2021; Martinez et al. 2023; Shikano et al. 2018b).

Mature human FAM237B and FAM237A share considerable sequence similarity, especially at their C-terminal fragment (Fig 1a). Predicted by the AlphaFold algorithm, they also displayed similar three-dimensional structures, with their C-terminal fragment forming a long α-helix (Fig 1b). A previous study reported that FAM237B is a weak agonist for GPR83: as high as 10 nM of FAM237B had no detectable activation effect in an inositol 1-phosphate accumulation assay (Sallee et al. 2020). In the present study, we prepared mature human FAM237B via an intein-fusion approach and measured its activity towards human GPR83 using NanoBiT-based ligand−receptor binding assay and NanoBiT-based β-arrestin recruitment assay. The results showed that FAM237B indeed displayed lower activity than its paralog FAM237A in both binding and activation assays; however, it could cause significant activation effect on human GPR83 at the nanomolar range (1□10 nM). Thus, FAM237B might be another endogenous agonist for GPR83.

**Fig. 1.**
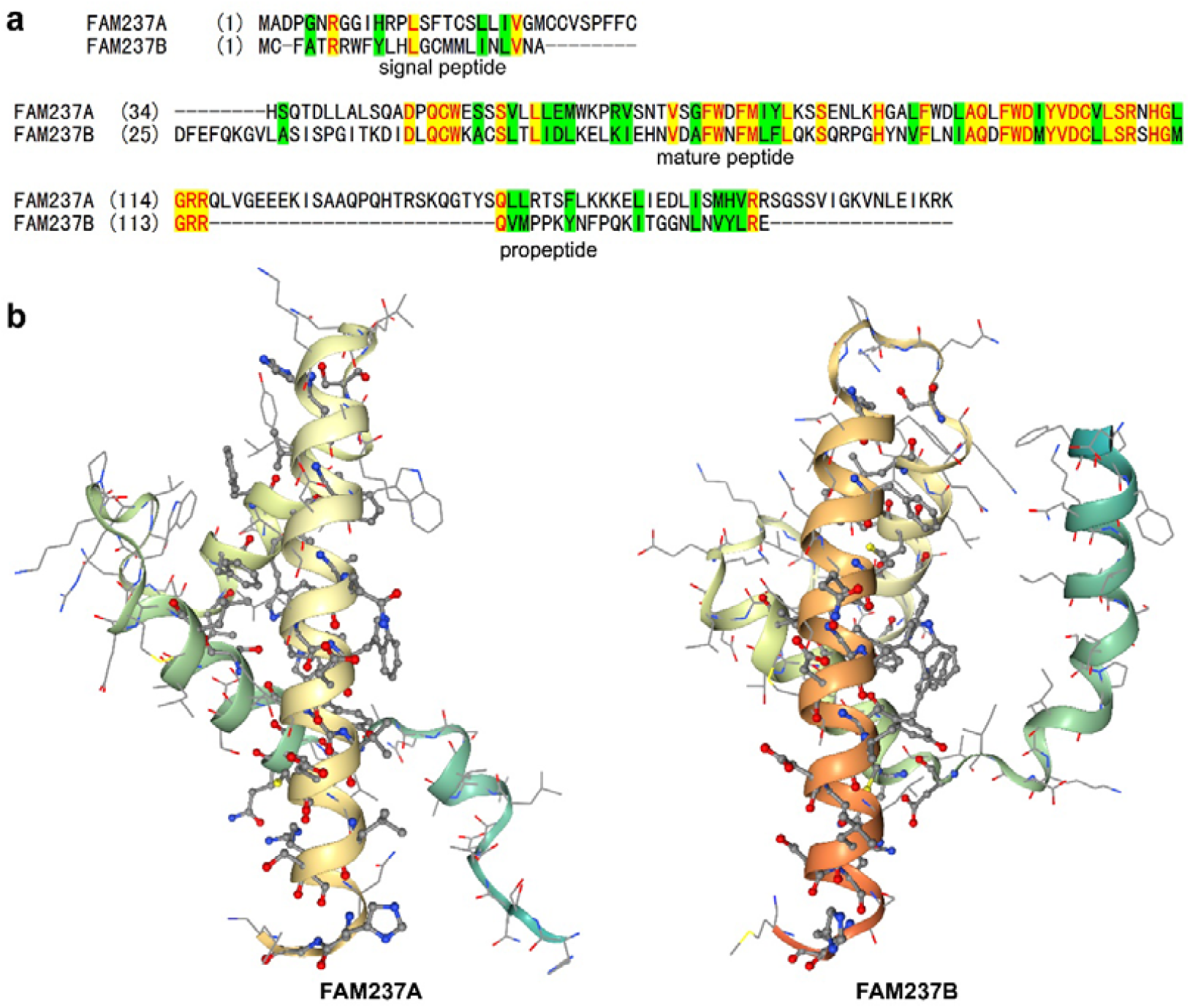
Comparison of human FAM237A and FAM237B. (**a**) Amino acid sequence alignment of the precursors of human FAM237A and FAM237B. (**b**) Three-dimensional structures of mature FAM237A (AF-A0A1B0GTK4-F1-model_v3) and FAM237B (AF-A0A1B0GVD1-F1-model_v3) predicted by AlphaFold algorithm. The residues identical in FAM237A and FAM237B are shown as sticks-and-balls, others are shown as lines.

## Materials and methods

### Preparation of mature FAM237B and the SmBiT-fused version via an intein-fusion approach

The nucleotide sequence encoding the C-terminally intein-fused mature human FAM237B (FAM237B-Intein) was chemically synthesized at Tsingke Biotechnology (Beijing, China). After cleavage with restriction enzymes NsiI and NotI, the synthetic DNA fragment was ligated into a pET vector, resulting in the bacterial overexpression construct pET/FAM237B-Intein-6×His (Fig. S2). The expression construct for the N-terminally SmBiT-fused FAM237B-Intein (SmBiT-FAM237B-Intein) was generated by polymerase chain reaction using pET/FAM237B-Intein-6×His as a template. After cleavage with NsiI and NotI, the amplified DNA fragment was ligated into the pET vector, resulting in the construct pET/SmBiT-FAM237B-Intein-6×His (Fig. S2). Their nucleotide sequence was confirmed by DNA sequencing.

Preparation of the mature FAM237B and the N-terminally SmBiT-fused FAM237B (SmBiT-FAM237B) was according to our previous procedure developed for preparation of mature FAM237A (Li et al. 2023). Briefly, the intein-fused precursor was solubilized from inclusion bodies via an *S*-sulfonation approach after overexpression in the *Escherichia coli* strain BL21(DE3), purified by immobilized metal ion affinity chromatography (Ni^2+^ column), and then treated by tris(2-carboxyethyl)phosphine and sodium 2-mercaptoethanesulfonate under denatured condition to remove the reversibly modified sulfonate moieties. Thereafter, the fully reduced precursor was subjected to renaturation and self-cleavage to release FAM237B thioester intermediate. After ammoniation, the amidated FAM237B was collected from the pellet, solubilized via *S*-sulfonation, and applied to a Ni^2+^ column to absorb the un-cleaved precursor and the cleaved intein. The flow-through fraction was then applied to high performance liquid chromatography (HPLC) and the *S*-sulfonated FAM237B was eluted from a C_4_ reverse-phase column (Hi-Pore reversed-phase column, 4.6 × 250 mm, Bio-Rad, Hercules, CA, USA) by an acidic acetonitrile gradient. After lyophilization, the linear FAM237B was subjected to *in vitro* refolding and the refolded FAM237B was purified by HPLC using the C_4_> reverse-phase column. The acidic acetonitrile gradient is composed of solvent A (0.1% aqueous trifluoroacetic acid) and solvent B (acetonitrile containing 0.1% trifluoroacetic acid). The purified mature FAM237B and SmBiT-FAM237B were dissolved in 1.0 mM aqueous hydrochloride (pH3.0), quantified by ultraviolet absorbance at 280 nm, aliquoted, and stored at -80 ºC for later activity assays.

### The NanoBiT-based ligand−receptor binding assay

The NanoBiT-based binding assay was conducted according to our previous procedure (Li et al. 2023). Briefly, human embryonic kidney (HEK) 293T cells were transiently transfected with the expression construct pcDNA6/sLgBiT-GPR83 that encodes an N-terminally secretory large NanoLuc fragment (sLgBiT)-fused human GPR83. Next day of transfection, the cells were trypsinized, seeded into white a opaque 96-well plate, and continuously cultured for ∼24 h to ∼90% confluence. To conduct the binding assay, the medium was removed and the binding solution (serum-free DMEM plus 0.1% BSA and 0.01% Tween-20) was added (50 µl/well). For saturation binding assay, the binding solution contained varied concentrations of SmBiT-FAM237B or SmBiT-FAM237A. For competition binding assay, the binding solution contained a constant concentration of SmBiT-FAM237A and varied concentrations of competitor. After incubation at 22 °C for ∼30 min, diluted NanoLuc substrate was added (10 µl/well, 30-fold dilution using the binding solution) and bioluminescence was immediately measured on a SpectraMax iD3 plate reader (Molecular Devices, Sunnyvale, CA, USA). The measured binding data were expressed as mean ± standard deviation (SD) (*n* = 3), and fitted to one-site binding model using SigmaPlot 10.0 software (SYSTAT software, Chicago, IL, USA).

### NanoBiT-based β-arrestin recruitment assay

The NanoBiT-based β-arrestin recruitment assay was conducted according to our previous procedure (Li et al. 2023). Briefly, HEK293T cells were transiently transfected with the expression control vector pCMV-TRE3G (Clontech, Mountain View, CA, USA) and the coexpression construct pTRE3G-BI/GPR83-LgBiT:SmBiT-ARRB2 that encodes a C-terminally LgBiT-fused human GPR83 (LgBiT-GPR83) and an N-terminally SmBiT-fused human β-arrestin 2 (SmBiT-ARRB2) via an inducible bi-directional promoter. Next day, the transfected cells were trypsinized, suspended in the induction medium (complete DMEM medium plus 2.5 ng/ml of doxycycline), seeded into a white opaque 96-well plate, and continuously cultured at 37 °C for ∼24 h. Before the activation assay, the cells were placed at room temperature for ∼30 min. To conduct the activation assay, the medium was removed and pre-warmed activation solution (serum-free DMEM plus 1% BSA) containing NanoLuc substrate was added to the living cells (40 µl/well, containing 1.0 µl of NanoLuc substrate stock). Thereafter, bioluminescence data were immediately collected for ∼4 min on a SpectraMax iD3 plate reader (Molecular Devices) with an interval of 11 s. Subsequently, peptide solution (diluted in the activation solution) was added to these wells (10 µl/well), and bioluminescence data were continuously collected for ∼10 min with an interval of 11 s. The measured absolute signals were corrected for inter well variability by forcing all curves after addition of NanoLuc substrate (without ligand) to same level.

## Results and discussion

### Preparation of mature FAM237B via an intein-fusion approach

To prepare mature FAM237B with the correct posttranslational modifications, we employed an intein-fusion approach, which was used to prepare active FAM237A in our recent study (Li et al. 2023). After overexpression in *E. coli*, the C-terminally intein-fused FAM237B precursor formed inclusion bodies, as analyzed by SDS-PAGE (Fig. 2a). After solubilized from the inclusion bodies using an *S*-sulfonation approach, the FAM237B precursor was purified using immobilized metal ion affinity chromatography, and then subjected to *in vitro* renaturation and self-cleavage. As analyzed by SDS-PAGE (Fig. 2a), approximately 70% of the fusion protein underwent self-cleavage, resulting in a weak band (indicated using an asterisk), presumably the released FAM237B thioester intermediate, at the bottom of the gel.

**Fig. 2.**
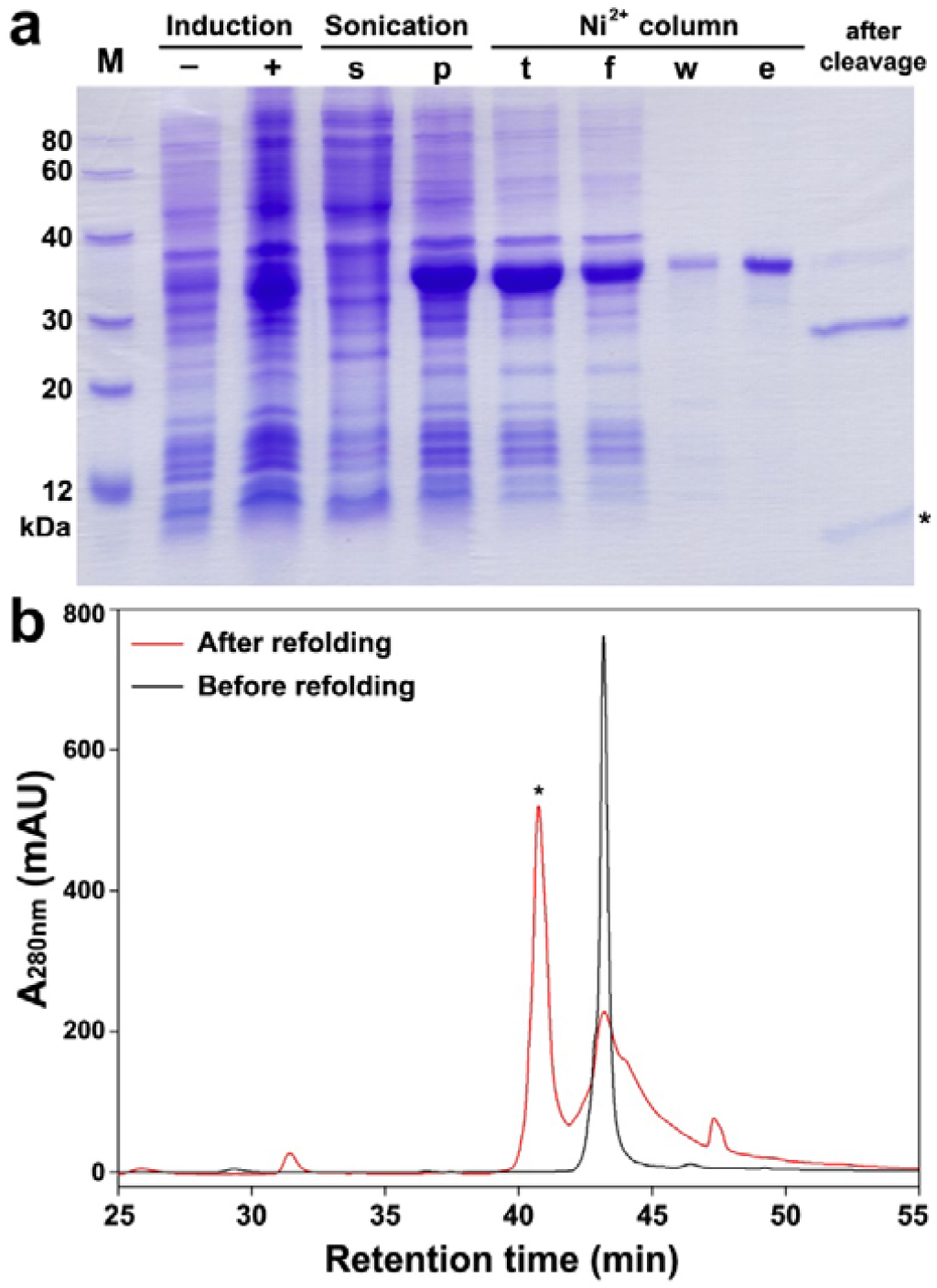
Preparation of mature FAM237B via an intein-fusion approach. (**a**) SDS-PAGE analysis of the intein-fused FAM237B precursor at different preparation steps. Lane (−), before IPTG induction; lane (+), after IPTG induction; lane (s), supernatant after sonication; lane (p), pellet after sonication; lane (t), the *S*-sulfonated fraction loading to the Ni^2+^ column; lane (f), flow-through fraction; lane (w), washing fraction by 30 mM imidazole; lane (e), eluted fraction by 250 mM imidazole; lane (after cleavage), after renaturation and overnight self-cleavage. Samples at different preparation stages were loaded onto a 15% SDS-gel, and the gel was stained by Coomassie brilliant blue R250 after electrophoresis. The band of the released FAM237B thioester intermediate is indicated by an asterisk. (**b**) HPLC analysis of the *S*-sulfonated FAM237B before and after *in vitro* refolding. ∼100 µg of sample before or after refolding was loaded onto a C_4_ reverse-phase column and eluted by an acidic acetonitrile gradient. The peak of the refolded FAM237B is indicated by an asterisk.

Thereafter, the released FAM237B thioester intermediate was ammoniated, and the α-amidated FAM237B linear peptide was obtained after several preparation steps, as analyzed using HPLC with a C_4_ reverse-phase column (Fig. 2b). After *in vitro* refolding, a new peak (indicated using an asterisk) with a shorter retention time was eluted from the C_4_ reverse-phase column (Fig. 2b), suggesting formation of the intramolecular disulfide bond. As estimated from its peak area, the yield of refolded FAM237B was ∼60%. From 1.0 liter of *E. coli* culture broth, ∼0.5 mg of mature FAM237B could be typically obtained.

### Mature FAM237B binds to GPR83 with moderately lower potency

We measured the binding of FAM237B to GPR83 using the NanoBiT-based binding assay, which relies on an N-terminally sLgBiT-fused receptor and an SmBiT-tagged ligand. After the recombinant SmBiT-FAM237B was added to living HEK293T cells overexpressing sLgBiT-GPR83, the measured bioluminescence increased in a saturable manner as the tracer concentration increased (Fig. 3a). After competition using 1.0 μM of mature FAM237A, the measured bioluminescence decreased markedly (Fig. 3a), suggesting that SmBiT-FAM237B can bind to sLgBiT-GPR83 and induce complementation of the ligand-fused SmBiT with the receptor-fused sLgBiT because of a proximity effect. Calculated from the saturation binding curve, the dissociation constant (K_d_) of SmBiT-FAM237B with sLgBiT-GPR83 was ∼300 nM. As a positive control, SmBiT-FAM237A bound to sLgBiT-GPR83 with a calculated K_d_ value of ∼80 nM (Fig. 3a). Thus, it seemed that SmBiT-FAM237B has a moderately lower binding affinity towards sLgBiT-GPR83 than SmBiT-FAM237A.

**Fig. 3.**
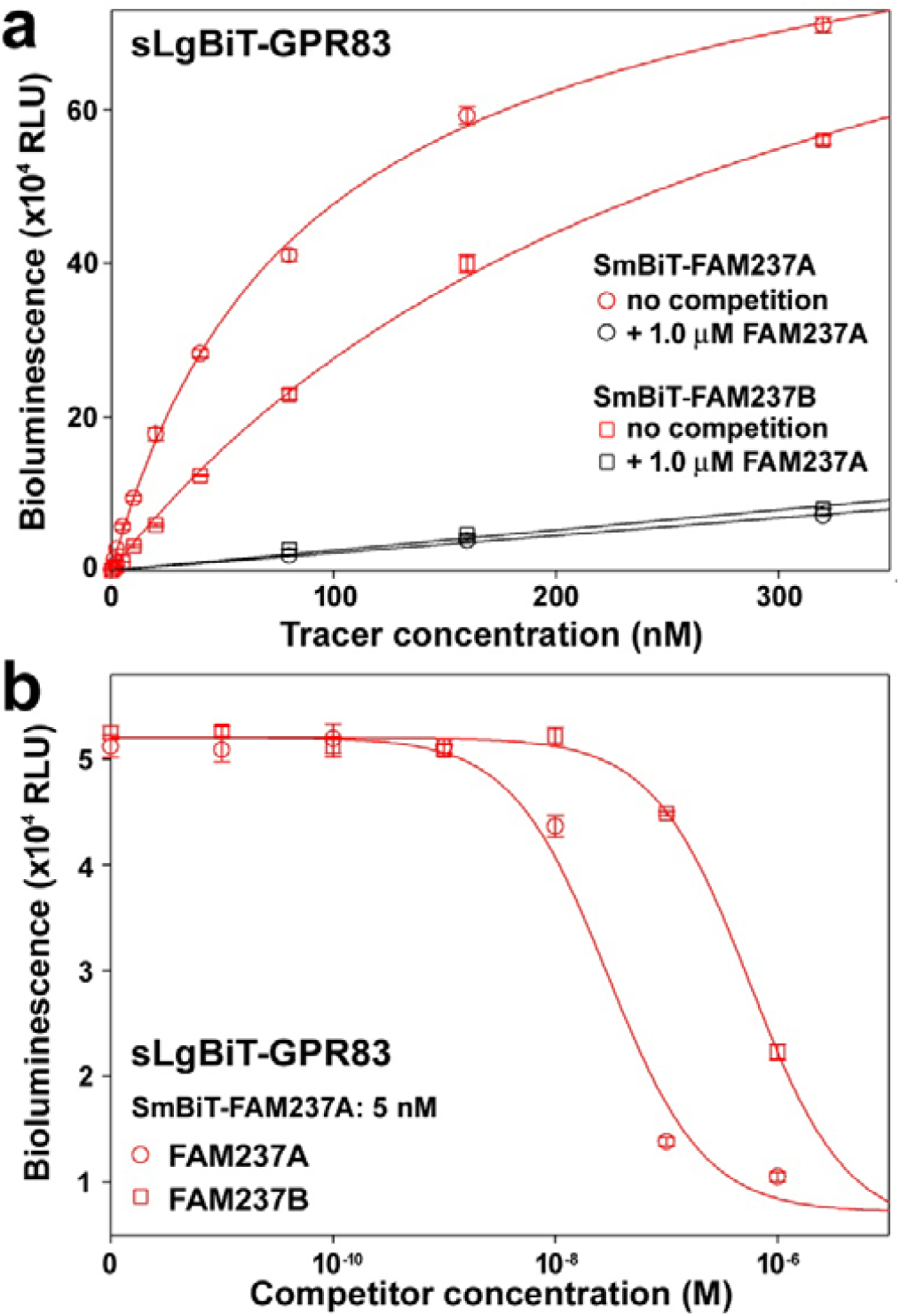
Binding of FAM237B with receptor GPR83 measured using the NanoBiT-based binding assay. (**a**) Saturation binding of SmBiT-FAM237B with HEK293T cells overexpressing sLgBiT-GPR83. SmBiT-FAM237A was used as a positive control. (**b**) Competition binding of mature FAM237B and FAM237A with sLgBiT-GPR83 using SmBiT-FAM237A as a tracer. The measured binding data are expressed as mean ± SD (*n* = 3) and fitted to one-site binding model using the SigmaPlot10.0 software.

Subsequently, we measured the binding potency of mature FAM237B with receptor GPR83 via competition binding assay using SmBiT-FAM237A as a tracer (Fig. 3b). Compared with mature FAM237A, mature FAM237B displayed approximately 20-fold lower receptor binding potency. Thus, it appeared that mature FAM237B has moderately lower binding potency with receptor GPR83 compared with its paralog FAM237A.

### Mature FAM237B activates GPR83 with moderately lower efficiency

We measured the activation effect of FAM237B towards GPR83 using the NanoBiT-based β-arrestin recruitment assay. As a positive control, mature FAM237A activated GPR83 efficiently: as low as 0.1 nM of FAM237A could cause a significant activation effect (Fig. 4a). For mature FAM237B, 0.1 nM had no detectable effect; 1.0 nM had a detectable but weak effect; and 10 nM caused a significant activation effect similar to that caused by 1.0 nM of FAM237A (Fig. 4b). Thus, FAM237B in the nanomolar range (1□10 nM) could cause a significant activation effect on GPR83, although its activation potency is lower than that of its paralog FAM237A.

**Fig. 4.**
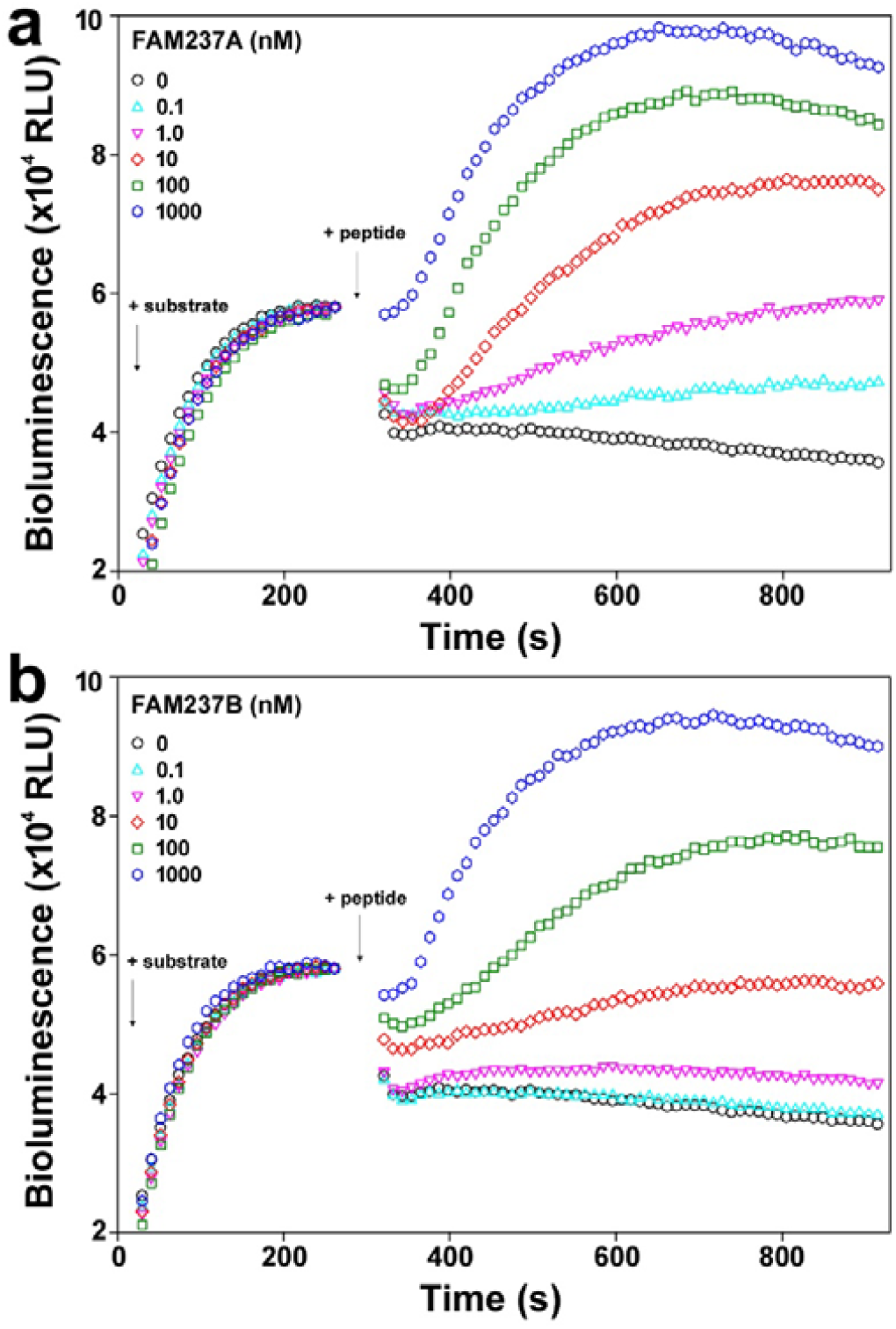
Activation of GPR83 by different concentrations of mature FAM237A (**a**) and FAM237B (**b**) measured by the NanoBiT-based β-arrestin recruitment assay. The measured bioluminescence data after sequential addition of NanoLuc substrate and peptide were plotted versus time using the SigmaPlot10.0 software.

To date, neuropeptide FAM237B has rarely been studied, although preliminary studies suggested that it is an orexigenic peptide implicated in the regulation of food intake and fat accumulation (Kato et al. 2021; Martinez et al. 2023; Shikano et al. 2018b). A previous study demonstrated that FAM237B can activate receptor GPR83 in an inositol 1-phosphate accumulation assay; however, its potency was much lower (∼100-fold) than that of FAM237A (Sallee et al. 2020) in this assay: 10 nM of FAM237B had no detectable activation effect (Sallee et al. 2020). In the present study, we demonstrated that in the nanomolar range (1□10 nM), FAM237B can cause a significant activation effect on GPR83 according to the NanoBiT-based β-arrestin recruitment assay. FAM237B also retains considerable binding affinity towards GPR83 compared with its paralog FAM237A. Thus, it seemed that FAM237B might be another endogenous agonist of GPR83. Further studies are needed to clarify whether GPR83 mediates the function of FAM237B *in vivo*.

Preparation of mature FAM237B is quite difficult because of its complex post-translational modifications and hydrophobic nature. Our present study provided an efficient approach to prepare active FAM237B via bacterial overexpression of a C-terminally intein-fused precursor and subsequent processing. In future, this approach could be used by other laboratories to prepare highly active FAM237B peptide for further functional studies.

## Abbreviations

BSA: Bovine serum albumin
FAM237A: Family with sequence similarity 237 member A
FAM237B: Family with sequence similarity 237 member B
GPR83: Gprotein-coupled receptor 83
HPLC: High performance liquid chromatography
NanoBiT: NanoLuc Binary Technology
PCSK1N: Proprotein convertase subtilisin/kexin type 1 inhibitor
SDS-PAGE: Sodium dodecyl sulfate-polyacrylamide gel electrophoresis
sLgBiT: Secretory large NanoLuc fragment
SmBiT: Low-affinity small complementation tag
TMD: Transmembrane domain

## Author Contributions

HZL, YFW, and WFH conducted the experiments; YLL and ZGX analyzed the data; ZYG conceived the research and wrote the manuscript; HZL and YFW contributed equally to this work. All authors have read and agreed to the published version of the manuscript.

## Funding

This work was supported by grant from the National Natural Science Foundation of China (grant no. 31971193).

## Data availability

The data that support the findings of this study are available from the corresponding author upon request.

## Declarations

### Competing Interests

The authors have no relevant financial or non-financial interests to disclose.

## Notes

### Competing Interest Statement

The authors have declared no competing interest.

