## Supplementary material for "Nanomolar range of FAM237B can activate receptor GPR83": Fig. S1-S2

### Contents:

**Fig. S1**. Amino acid sequence alignment of FAM237B orthologs from fishes to mammals.

**Fig. S2.** The nucleotide and amino acid sequences of the intein-fused human FAM237B precursors overexpressed in *E. coli*.

Amino acid sequence alignment of FAM237B orthologs from fishes to mammals

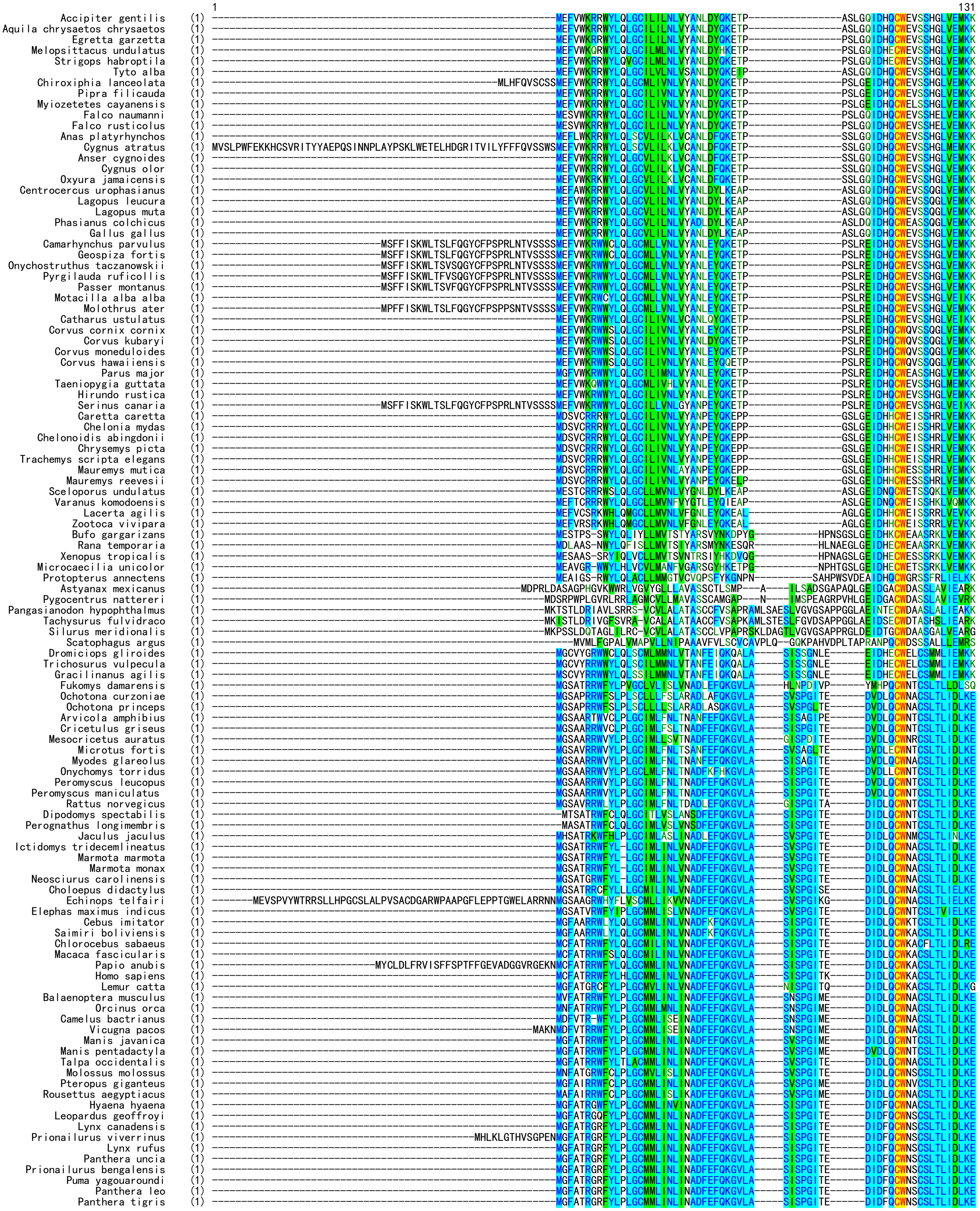

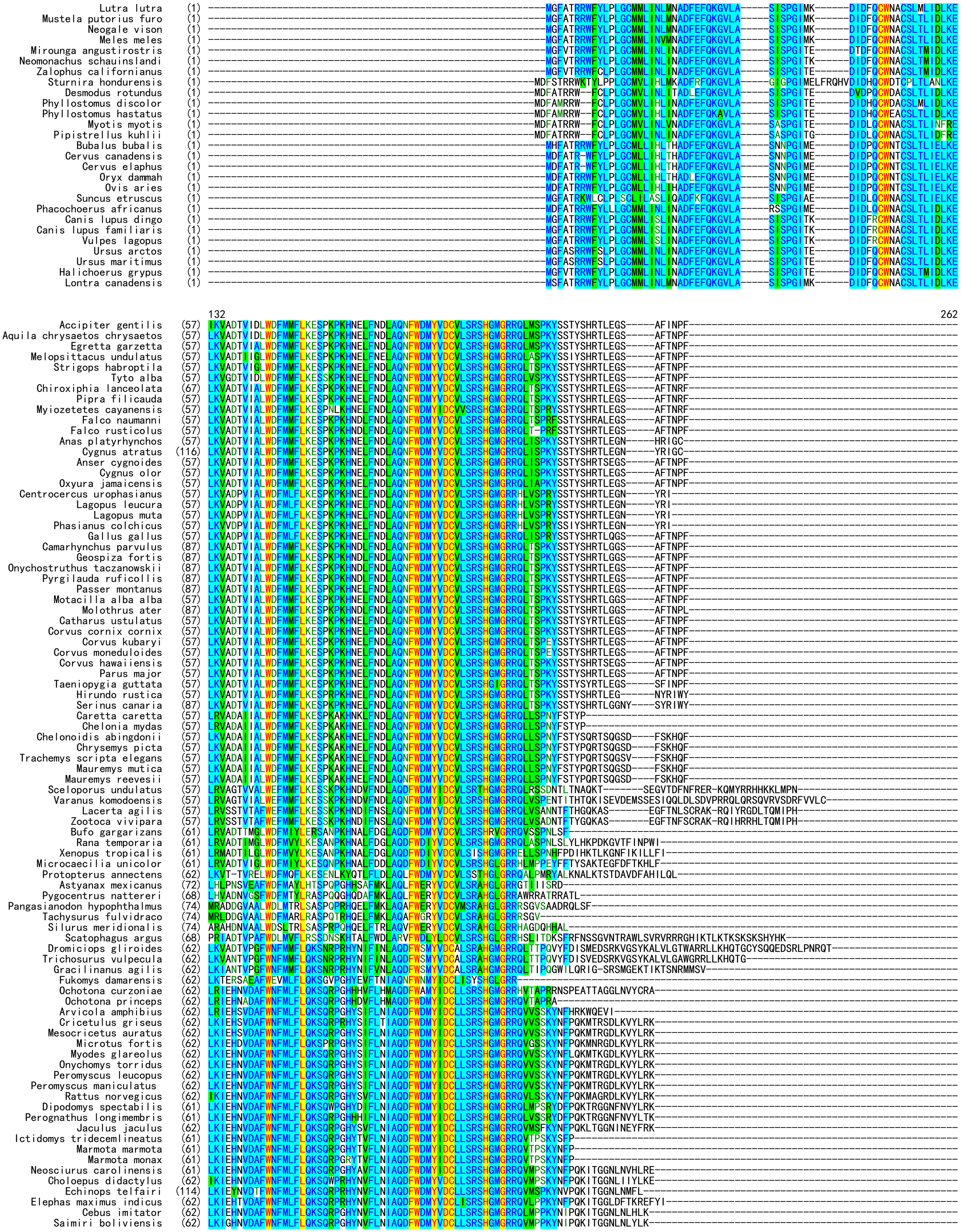

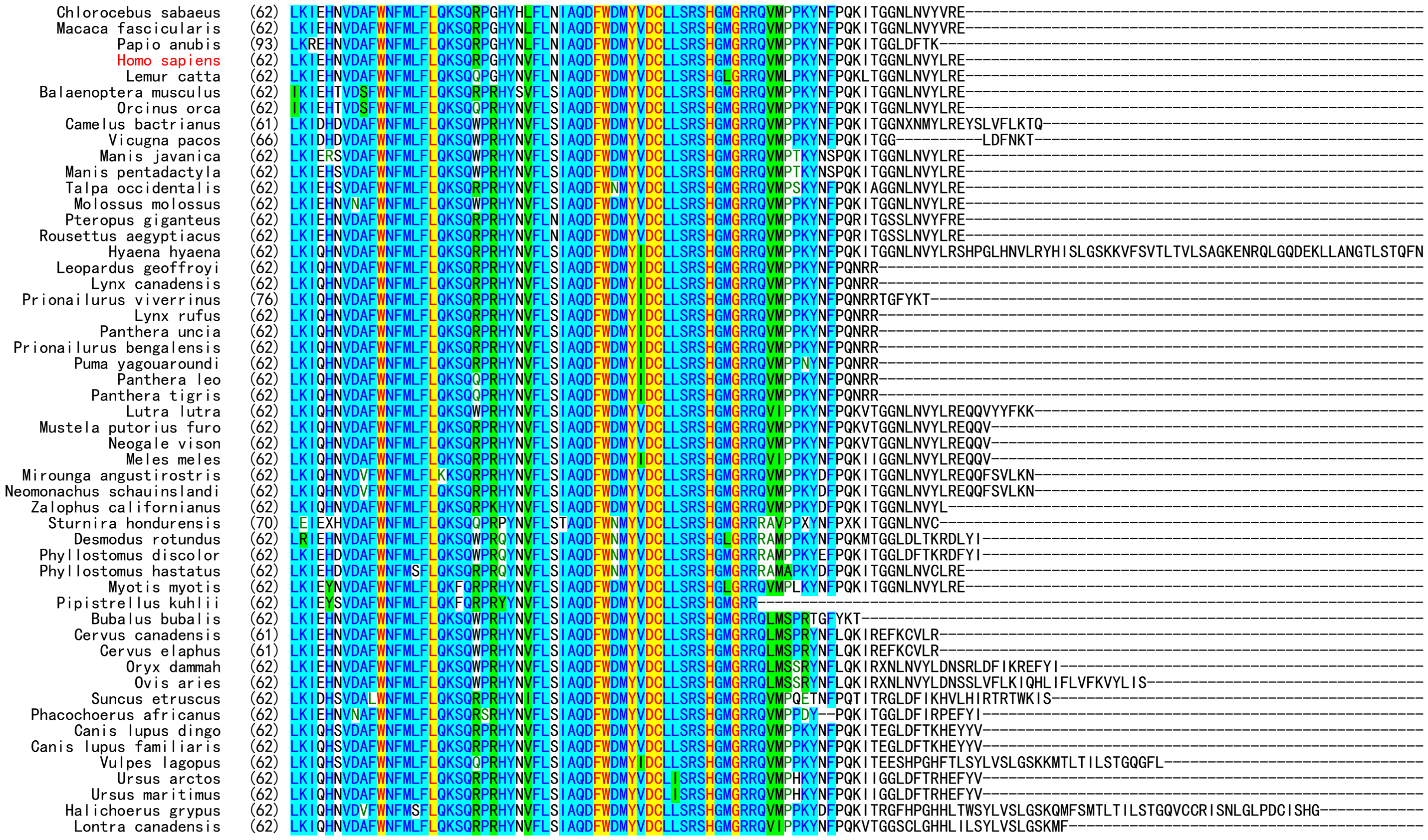


**Fig. S1.** Amino acid sequence alignment of FAM237B orthologs from fishes to mammals.

### FAM237B-Intein-6×His in pET vector

```
      NsiI
1     ATG CAT GAC TTT GAA TTT CAA AAA GGT GTG CTG GCC AGC ATC AGC CCG GGC ATC ACC AAA GAC ATT GAT CTC CAG
      TAC GTA CTG AAA CTT AAA GTT TTT CCA CAC GAC CGG TCG TAG TCG GGC CCG TAG TGG TTT CTG TAA CTA GAG GTC
       M   H   D   F   E   F   Q   K   G   V   L   A   S   I   S   P   G   I   T   K   D   I   D   L   Q

76    TGC TGG AAA GCT TGT TCT TTG ACG TTA ATT GAT CTG AAG GAA CTG AAG ATC GAG CAC AAT GTG GAC GCG TTT TGG
      ACG ACC TTT CGA ACA AGA AAC TGC AAT TAA CTA GAC TTC CTT GAC TTC TAG CTC GTG TTA CAC CTG CGC AAA ACC
       C   W   K   A   C   S   L   T   L   I   D   L   K   E   L   K   I   E   H   N   V   D   A   F   W

151   AAC TTC ATG TTG TTC TTA CAA AAA TCC CAG CGC CCT GGT CAT TAC AAT GTC TTC CTG AAC ATT GCG CAG GAT TTC
      TTG AAG TAC AAC AAG AAT GTT TTT AGG GTC GCG GGA CCA GTA ATG TTA CAG AAG GAC TTG TAA CGC GTC CTA AAG
       N   F   M   L   F   L   Q   K   S   Q   R   P   G   H   Y   N   V   F   L   N   I   A   Q   D   F

226   TGG GAC ATG TAT GTT GAC TGC CTG CTT TCA CGT AGT CAT GGC ATG TGT ATT TGC GGT GAT GCT CTG GTG GCC CTG
      ACC CTG TAC ATA CAA CTG ACG GAC GAA AGT GCA TCA GTA CCG TAC ACA TAA ACG CCA CTA CGA GAC CAC CGG GAC
       W   D   M   Y   V   D   C   L   L   S   R   S   H   G   M   C   I   C   G   D   A   L   V   A   L

301   CCG GAA GGC GAA AGC GTT CGT ATC GCA GAC ATT GTC CCA GGT GCT CGC CCG AAC AGT GAT AAT GCG ATC GAC CTG
      GGC CTT CCG CTT TCG CAA GCA TAG CGT CTG TAA CAG GGT CCA CGA GCG GGC TTG TCA CTA TTA CGC TAG CTG GAC
       P   E   G   E   S   V   R   I   A   D   I   V   P   G   A   R   P   N   S   D   N   A   I   D   L

376   AAA GTA CTC GAT CGT CAC GGC AAC CCA GTG TTA GCG GAT CGT TTG TTT CAT TCT GGT GAG CAT CCT GTT TAT ACC
      TTT CAT GAG CTA GCA GTG CCG TTG GGT CAC AAT CGC CTA GCA AAC AAA GTA AGA CCA CTC GTA GGA CAA ATA TGG
       K   V   L   D   R   H   G   N   P   V   L   A   D   R   L   F   H   S   G   E   H   P   V   Y   T

451   GTC CGC ACG GTA GAA GGT CTG CGT GTG ACT GGC ACC GCC AAC CAC CCG CTG CTG TGC TTA GTT GAT GTC GCA GGT
      CAG GCG TGC CAT CTT CCA GAC GCA CAC TGA CCG TGG CGG TTG GTG GGC GAC GAC ACG AAT CAA CTA CAG CGT CCA
       V   R   T   V   E   G   L   R   V   T   G   T   A   N   H   P   L   L   C   L   V   D   V   A   G

526   GTA CCG ACC CTG TTG TGG AAG TTA ATT GAT GAG ATC AAA CCA GGC GAC TAC GCT GTG ATT CAG CGT TCC GCG TTC
      CAT GGC TGG GAC AAC ACC TTC AAT TAA CTA CTC TAG TTT GGT CCG CTG ATG CGA CAC TAA GTC GCA AGG CGC AAG
       V   P   T   L   L   W   K   L   I   D   E   I   K   P   G   D   Y   A   V   I   Q   R   S   A   F

601   TCA GTT GAT TGT GCT GGC TTT GCC CGC GGT AAA CCG GAG TTC GCA CCT ACC ACT TAT ACG GTG GGC GTT CCG GGT
      AGT CAA CTA ACA CGA CCG AAA CGG GCG CCA TTT GGC CTC AAG CGT GGA TGG TGA ATA TGC CAC CCG CAA GGC CCA
       S   V   D   C   A   G   F   A   R   G   K   P   E   F   A   P   T   T   Y   T   V   G   V   P   G

676   CTG GTC CGT TTC CTG GAG GCG CAC CAT CGC GAT CCA GAC GCA CAG GCT ATC GCG GAT GAA CTG ACG GAT GGC CGT
      GAC CAG GCA AAG GAC CTC CGC GTG GTA GCG CTA GGT CTG CGT GTC CGA TAG CGC CTA CTT GAC TGC CTA CCG GCA
       L   V   R   F   L   E   A   H   H   R   D   P   D   A   Q   A   I   A   D   E   L   T   D   G   R

751   TTT TAT TAC GCC AAA GTG GCT AGC GTT ACC GAC GCG GGT GTG CAA CCG GTC TAT TCG TTA CGC GTT GAT ACG GCA
      AAA ATA ATG CGG TTT CAC CGA TCG CAA TGG CTG CGC CCA CAC GTT GGC CAG ATA AGC AAT GCG CAA CTA TGC CGT
       F   Y   Y   A   K   V   A   S   V   T   D   A   G   V   Q   P   V   Y   S   L   R   V   D   T   A

                                                          NotI
826   GAC CAC GCG TTC ATC ACG AAC GGC TTC GTT TCT CAT GCG GCG GCC GCA CTC GAG CAC CAC CAC CAC CAC CAC TGA
      CTG GTG CGC AAG TAG TGC TTG CCG AAG CAA AGA GTA CGC CGC CGG CGT GAG CTC GTG GTG GTG GTG GTG GTG ACT
       D   H   A   F   I   T   N   G   F   V   S   H   A   A   A   A   L   E   H   H   H   H   H   H   *
```

### SmBiT-FAM237B-Intein-6×His in pET vector

```
      NsiI
1     ATG CAT GTG ACC GGC TAC CGT CTG TTT GAA GAA ATT CTG GGC GGC AGC GGT GGT GGC GAC TTT GAA TTT CAA AAA
      TAC GTA CAC TGG CCG ATG GCA GAC AAA CTT CTT TAA GAC CCG CCG TCG CCA CCA CCG CTG AAA CTT AAA GTT TTT
       M   H   V   T   G   Y   R   L   F   E   E   I   L   G   G   S   G   G   G   D   F   E   F   Q   K

76    GGT GTG CTG GCC AGC ATC AGC CCG GGC ATC ACC AAA GAC ATT GAT CTC CAG TGC TGG AAA GCT TGT TCT TTG ACG
      CCA CAC GAC CGG TCG TAG TCG GGC CCG TAG TGG TTT CTG TAA CTA GAG GTC ACG ACC TTT CGA ACA AGA AAC TGC
       G   V   L   A   S   I   S   P   G   I   T   K   D   I   D   L   Q   C   W   K   A   C   S   L   T

151   TTA ATT GAT CTG AAG GAA CTG AAG ATC GAG CAC AAT GTG GAC GCG TTT TGG AAC TTC ATG TTG TTC TTA CAA AAA
      AAT TAA CTA GAC TTC CTT GAC TTC TAG CTC GTG TTA CAC CTG CGC AAA ACC TTG AAG TAC AAC AAG AAT GTT TTT
       L   I   D   L   K   E   L   K   I   E   H   N   V   D   A   F   W   N   F   M   L   F   L   Q   K

226   TCC CAG CGC CCT GGT CAT TAC AAT GTC TTC CTG AAC ATT GCG CAG GAT TTC TGG GAC ATG TAT GTT GAC TGC CTG
      AGG GTC GCG GGA CCA GTA ATG TTA CAG AAG GAC TTG TAA CGC GTC CTA AAG ACC CTG TAC ATA CAA CTG ACG GAC
       S   Q   R   P   G   H   Y   N   V   F   L   N   I   A   Q   D   F   W   D   M   Y   V   D   C   L

301   CTT TCA CGT AGT CAT GGC ATG TGT ATT TGC GGT GAT GCT CTG GTG GCC CTG CCG GAA GGC GAA AGC GTT CGT ATC
      GAA AGT GCA TCA GTA CCG TAC ACA TAA ACG CCA CTA CGA GAC CAC CGG GAC GGC CTT CCG CTT TCG CAA GCA TAG
       L   S   R   S   H   G   M   C   I   C   G   D   A   L   V   A   L   P   E   G   E   S   V   R   I

376   GCA GAC ATT GTC CCA GGT GCT CGC CCG AAC AGT GAT AAT GCG ATC GAC CTG AAA GTA CTC GAT CGT CAC GGC AAC
      CGT CTG TAA CAG GGT CCA CGA GCG GGC TTG TCA CTA TTA CGC TAG CTG GAC TTT CAT GAG CTA GCA GTG CCG TTG
       A   D   I   V   P   G   A   R   P   N   S   D   N   A   I   D   L   K   V   L   D   R   H   G   N

451   CCA GTG TTA GCG GAT CGT TTG TTT CAT TCT GGT GAG CAT CCT GTT TAT ACC GTC CGC ACG GTA GAA GGT CTG CGT
      GGT CAC AAT CGC CTA GCA AAC AAA GTA AGA CCA CTC GTA GGA CAA ATA TGG CAG GCG TGC CAT CTT CCA GAC GCA
       P   V   L   A   D   R   L   F   H   S   G   E   H   P   V   Y   T   V   R   T   V   E   G   L   R

526   GTG ACT GGC ACC GCC AAC CAC CCG CTG CTG TGC TTA GTT GAT GTC GCA GGT GTA CCG ACC CTG TTG TGG AAG TTA
      CAC TGA CCG TGG CGG TTG GTG GGC GAC GAC ACG AAT CAA CTA CAG CGT CCA CAT GGC TGG GAC AAC ACC TTC AAT
       V   T   G   T   A   N   H   P   L   L   C   L   V   D   V   A   G   V   P   T   L   W   K   L
```

```
                            NotI
```

```
601   ATT GAT GAG ATC AAA CCA GGC GAC TAC GCT GTG ATT CAG CGT TCC GCG TTC TCA GTT GAT TGT GCT GGC TTT GCC
      TAA CTA CTC TAG TTT GGT CCG CTG ATG CGA CAC TAA GTC GCA AGG CGC AAG AGT CAA CTA ACA CGA CCG AAA CGG
       I   D   E   I   K   P   G   D   Y   A   V   I   Q   R   S   A   F   S   V   D   C   A   G   F   A

676   CGC GGT AAA CCG GAG TTC GCA CCT ACC ACT TAT ACG GTG GGC GTT CCG GGT CTG GTC CGT TTC CTG GAG GCG CAC
      GCG CCA TTT GGC CTC AAG CGT GGA TGG TGA ATA TGC CAC CCG CAA GGC CCA GAC CAG GCA AAG GAC CTC CGC GTG
       R   G   K   P   E   F   A   P   T   T   Y   T   V   G   V   P   G   L   V   R   F   L   E   A   H

751   CAT CGC GAT CCA GAC GCA CAG GCT ATC GCG GAT GAA CTG ACG GAT GGC CGT TTT TAT TAC GCC AAA GTG GCT AGC
      GTA GCG CTA GGT CTG CGT GTC CGA TAG CGC CTA CTT GAC TGC CTA CCG GCA AAA ATA ATG CGG TTT CAC CGA TCG
       H   R   D   P   D   A   Q   A   I   A   D   E   L   T   D   G   R   F   Y   Y   A   K   V   A   S

826   GTT ACC GAC GCG GGT GTG CAA CCG GTC TAT TCG TTA CGC GTT GAT ACG GCA GAC CAC GCG TTC ATC ACG AAC GGC
      CAA TGG CTG CGC CCA CAC GTT GGC CAG ATA AGC AAT GCG CAA CTA TGC CGT CTG GTG CGC AAG TAG TGC TTG CCG
       V   T   D   A   G   V   Q   P   V   Y   S   L   R   V   D   T   A   D   H   A   F   I   T   N   G

                                 NotI
901   TTC GTT TCT CAT GCG GCG GCC GCA CTC GAG CAC CAC CAC CAC CAC CAC TGA
      AAG CAA AGA GTA CGC CGC CGG CGT GAG CTC GTG GTG GTG GTG GTG GTG ACT
       F   V   S   H   A   A   A   A   L   E   H   H   H   H   H   H   *
```

**Fig. S2.** The nucleotide and amino acid sequences of the intein-fused human FAM237B precursors overexpressed in *E. coli*. The amino acid sequence of mature human FAM237B is shown in red, and that of the intein in green, and that of SmBiT in blue.
